# Mosaic Patterns of B-vitamin Synthesis and Utilization in a Natural Marine Microbial Community

**DOI:** 10.1101/280438

**Authors:** Laura Gómez-Consarnau, Rohan Sachdeva, Scott M. Gifford, Lynda S. Cutter, Jed A. Fuhrman, Sergio A. Sañudo-Wilhelmy, Mary Ann Moran

## Abstract

Aquatic environments contain diverse microbial communities whose complex interactions mediate the cycling of major and trace nutrients such as vitamins. B-vitamins are essential coenzymes that many organisms cannot synthesize. Thus their exchange among de-novo synthesizers and auxotrophs is expected to play an important role in the microbial consortia and explain some of the temporal and spatial changes observed in diversity. In this study, we analyzed metatranscriptomes of a natural coastal microbial community, diel sampled-quarterly over one year to try to identify the potential major B-vitamin synthesizers and consumers. Our transcriptomic data show that the best-represented taxa dominated the expression of synthesis genes for some B-vitamins but lacked transcripts for others. For instance, Rhodobacterales dominated the expression of vitamin-B_12_ synthesis, but not of vitamin-B_7_, whose synthesis transcripts were mainly represented by Flavobacteria.In contrast, bacterial groups that constituted less than 4% of the community (e.g., Verrucomicrobia) accounted for most of the vitamin-B_1_ synthesis transcripts. Furthermore, ambient vitamin-B_1_ concentrations were higher in samples collected during the day, and were positively correlated with chlorophyll-*a* concentration. Our analysis supports the hypothesis that the mosaic of metabolic interdependencies through B-vitamin synthesis and exchange are key processes that contribute to shaping microbial communities in nature.

## Introduction

Aquatic environments contain large communities of microorganisms whose complex synergistic chemical interactions mediate the transfer of energy and the cycling of major and trace nutrients. Molecular studies have shown that marine microbial systems are extremely diverse, containing thousands of different taxa that combined presumably sustain ecosystem functioning (Sogin et al., 2006). In general, while a few species numerically dominate a particular microbial community, they are complemented by a large number of less abundant species whose specific function is not well understood (PedrÓs-AliÓ, 2012). For example, similar to the Hutchinson’s paradox of the plankton (Hutchinson, 1961), it is unclear why the marine heterotrophic bacterioplankton community is so diverse at certain locations if all share the same niche and ecological functions (Moran, 2008). Ultimately, recognizing the connectivity among the species that compose the heterotrophic bacterioplankton community represents a key to understanding the fate of up to half of the marine primary production that is routed through the microbial loop (Azam et al., 1983). Field and laboratory studies have shown that resource allocation from algal-derived organic matter could explain some marine bacterioplankton successions and the dominance of certain microbial taxa capable of distinct decomposition pathways (McCarren et al., 2010; Teeling et al., 2012). However, these mechanisms cannot explain the large background heterotrophic community that exists under non-bloom conditions during most of the year. In contrast, the bacterial connectivity and diversity observed in microbial communities could reflect complex interdependencies associated with external metabolites such as the B-vitamins, as observed in mixed culture experiments (Croft et al., 2005).

B-vitamins are the most versatile and ancient coenzymes (Monteverde et al., 2017). They catalyze a wide spectrum of critical metabolic reactions, such as the tricarboxylic acid (TCA) and Calvin cycles (thiamin; vitamin-B_1_; VB_1_), carbon fixation via the reverse TCA cycle (biotin; vitamin-B_7_; VB_7_) or the synthesis of methionine (cobalamin; vitamin-B_12_, VB_12_) (Sañudo-Wilhelmy et al., 2014). Vitamin B_6_ (pyridoxine, VB_6_) seems to be remarkably important as it catalyzes almost 2% of all prokaryotic functions (Percudani and Peracchi 2003) and yet, has rarely been studied in marine systems (Sañudo-Wilhelmy et al., 2012). Despite their relevance, B-vitamins auxotrophy is widespread among marine eukaryotes (Croft et al., 2006, Tang et al., 2010; Paerl et al., 2015) and may influence the taxonomic composition of phytoplankton communities as well as the rates of primary production and carbon fixation (Panzeca et al., 2006; Sañudo-Wilhelmy et al., 2006; Koch et al., 2011). For decades, it was assumed that marine prokaryotes were the source of exogenous B-vitamins for marine phytoplankton (Kurata 1986; Croft et al., 2005). However, the extensive genomic information that is now available suggests that not all bacteria and archaea are *de-novo* synthesizers (LeBlanc et al. 2011; Sañudo-Wilhelmy et al., 2014). For example, the highly streamlined *Candidatus* Pelagibacter ubique lacks the biosynthetic pathways for VB, VB, and VB, and can reach concentrations of 10^9^ cells ml^-1^ only when these vitamins are present, regardless of how much C, N or P is available (Carini et al., 2012). This example alone suggests that exogenous sources of B-vitamins (dissolved and/or particulate) must be available within the water column. The hypothetical direct or indirect “metabolic cooperation” (Raes and Bork, 2008) among the bacterio-and phytoplankton particle-associated communities via vitamin exchange has been intuited from results of experiments using isolated strains in culture (Croft et al., 2005; Wagner-Döbler et al., 2010; Kazamia et al., 2012). Furthermore, dissolved vitamins are also released into the environment through processes that do not involve direct microbial interactions, as VB_12_ excretion during cell division has been observed in pure cultures of cyanobacteria (Bonnet et al., 2010). Regardless of whether or not B-vitamin auxotrophs rely on direct or indirect microbial exchange to obtain these metabolites, the identification of the potential vitamin producers and consumers in a natural microbial community is important for understanding the processes sustaining the ecosystem. Recent work of Heal et al. (2017) studied the potential VB_12_ interdependencies among producers and consumers in the North Pacific Ocean. However, the study of these ectocrine interactions remains largely unfinished, as they have not been examined for other B-vitamins and in other marine environments.

In this study, we used metatranscriptomics to identify the transcriptional investment of different taxa in B-vitamin synthesis and utilization for VB_1_, VB_6_, VB_7_ and VB_12_ within a natural coastal marine microbial community. Our data show that some of the best-represented taxa dominated the expression of synthesis genes for some, but not all B-vitamins, suggesting the need for metabolite exchange among microbial groups. The concentration of dissolved VB_1_ and VB_6_ in this environment followed diel oscillations, with higher levels during the day compared to night. These environmental diel fluctuations of B-vitamins could be related to their metabolic function as protectants against oxidative stress and potentially constitute a mechanism of vitamin supply to the auxotrophic members of the community.

## Results and Discussion

### Microbial community composition

Taxonomic diversity of metabolically active bacteria through the different seasons was estimated using the total number of transcripts recruited for each microbial group (Gifford et al., 2014). Total community expression was represented by bacterial taxa that are typically found in marine coastal ecosystems (Gifford et al., 2013; 2014), including Alphaproteobacteria (Rhodobacterales, relative expression 11-18%; SAR11, 1-14%; SAR116, 3-7%),Flavobacteria (2-11%), Gammaproteobacteria (unclassified, 6-11%) and Betaproteobacteria (3-7%) (Figure 1 and Table S3). Eukaryotic mRNAs were the second-most expressed transcripts detected after Rhodobacterales in this 3.0-micron pre-filtered community (on average ∼11% of reads; range 4-38%), and were particularly important during periods of high chlorophyll-a (Chl-*a*) (Summer ‘08, Fall ‘08 and Spring ‘09, accounting for 18-38% of the total mRNA library). The microbial picoeukaryotic taxa with the highest expression in this 0.2 to 3.0 µm plankton size fraction were Micromonas (20-44% of eukaryotic sequences), Ostreococcus (20-39%) and Chlorella-like (5-36%; Table 2, Table S3). The relative increase in expression of certain bacterioplankton groups during the Chl-a peaks (e.g. Flavobacteria during the phytoplankton bloom of Winter ‘09) is consistent with similar trends repeatedly reported in the literature (Kirchman, 2002; Teeling et al., 2012). However, the relative contribution of the major microbial community taxa to total community transcripts remained relatively stable during our year-long study (e.g. SAR11, SAR116, Rhodobacterales, Betaproteobacteria; Figure 1, Table S3). Archaeal transcripts7.5-10.5% of all transcripts during summer 08 and were also high but to a lesser extent in summer ‘09 (2-4% of all transcripts), while they were low in all other seasons (Hollibaugh et al., 2011; 2013). Other taxonomic groups such as Cyanobacteria and Verrucomicrobia were also present throughout the year although comprising less than 4% of the prokaryotic expression (Figure 1, Table S3).

**Figure 1.**
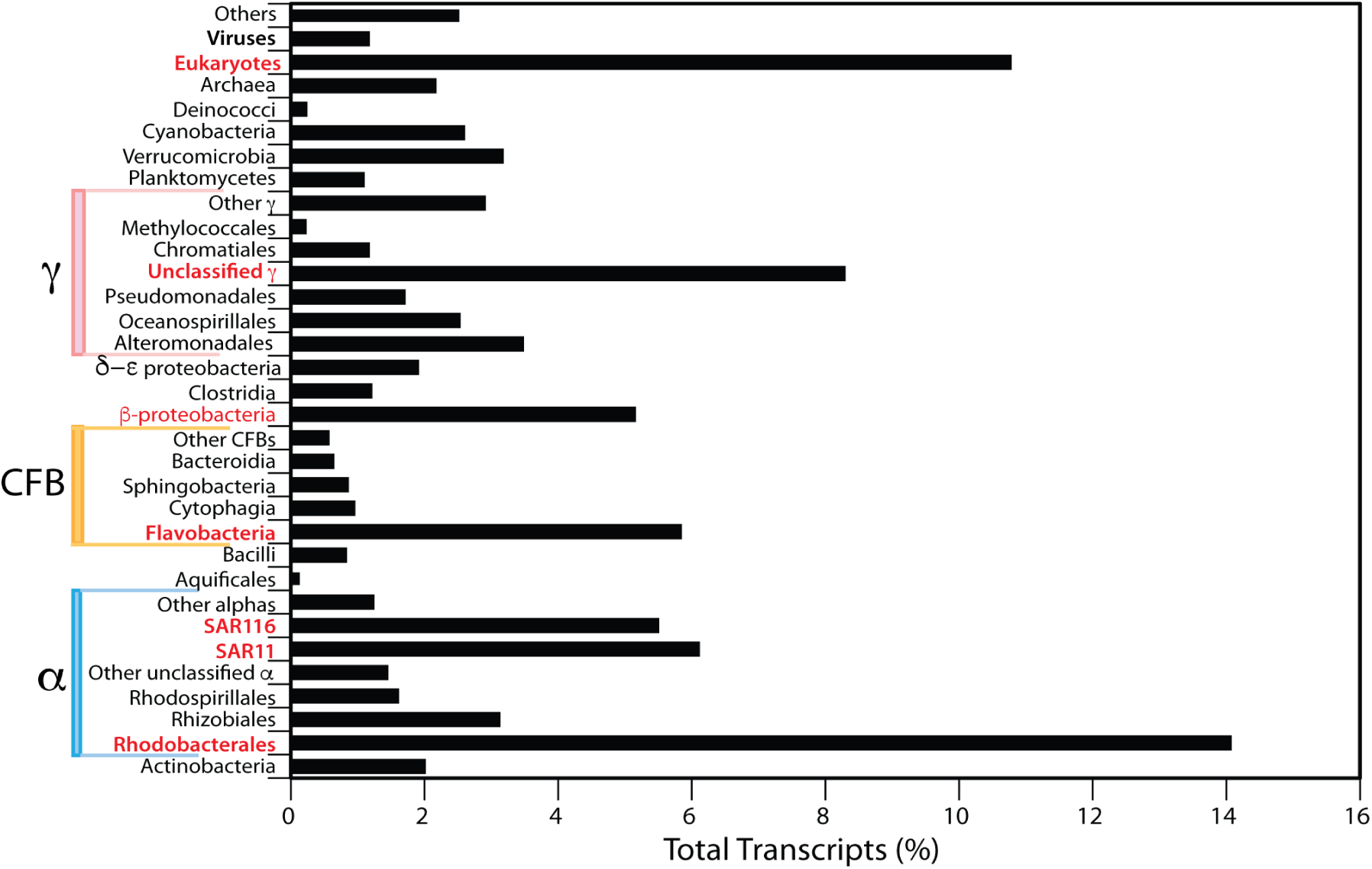
Percentage of the total transcripts detected in 16 southeastern U.S. metatranscriptomes, classified by taxon. Red font indicates the best-represented taxonomic groups as determined from mRNA abundance. The symbols α β γ δ and ε refer to the different proteobacteria classes. CFB refer to the Cytophaga-Flavobacterium-Bacteroides (CFB) phylum.

### Expression of vitamin synthesis and vitamin-dependent metabolisms

The correlation between gene expression and the actual associated metabolic process in the environment is not always easy to discern (Gifford et al., 2011). It is currently known that the ratio between mRNA and protein can be quite variable and may not quantitatively reflect the immediate cell response to environmental changes (Taniguchi et al., 2010). The inability to predict protein levels from mRNA could be explained by the long half-life of proteins compared to mRNA, in addition to variable post-transcriptional modifications and translation efficiencies among other factors (Moran et al., 2013). Nevertheless, metatranscriptomics is a powerful tool to identify patterns that can be used to identify the potential organisms involved in different environmental processes. To try to understand B-vitamin synthesis and utilization in this coastal environment, we quantified the total number of annotatable transcripts involved in accounting for almost 50% of all the environmental VB_12_ synthesis transcripts in some samples (VB_12_ synthesis:total transcript ratio had a residual of 27% from the community-wide vitamin regression line; Figure 2A, Table S4). This is in contrast to what was reported from incubation experiments in the Southern Ocean, in which Oceanospirillaceae appeared to be the major VB_12_ synthesizers (Bertrand et al., 2015). However, because Rhodobacterales VB_12_ dependence transcript abundance was also high, their VB_12_ dependence:synthesis transcript ratio was typical for this community (VB_12_ dependence:synthesis ratio had a residual of less than 2% from the community-wide regression line; Figure 2C). For other B-vitamins, though, Rhodobacterales vitamin-synthesis transcripts were clearly below the typical dependence:synthesis transcript relationship. For VB_1_ and VB_6_, only 7% and 12% of the synthesis transcripts belonged to Rhodobacterales, while they were responsible for a 2-fold higher percentage of the vitamin-dependent gene transcripts (dependence:synthesis residuals of 9% and 10%, respectively; Figure 2, Table S4). Their dependence:synthesis transcript relationship for VB_7_ was even more unbalanced with 2% of the community synthesis genes compared to 20% of the community requirement (synthesis-dependence residuals of 18%; Figure 2C, Table S3, S4). These field-based results are consistent with genomes from isolated marine strains showing that while almost all of the cultured Rhodobacterales can synthesize vitamin-B, only 50% and 70% can produce VB the synthesis of VB_1_, VB_6_, VB_7_ and VB^12^, as well as their and VB_7_ respectively (Sañudo-Wilhelmy et al., 2014). Figure vitamin-dependent enzymes (Table 1, Table S1). This semi-quantitative approach allows the evaluation of the overall transcript investment in B-vitamin synthesis and dependent functions within metatranscriptomes. On the synthesis side, the highest number of transcripts was found for VB_6_ synthesis (0.05% of the community transcriptome), followed by VB_12_,VB_1_, and VB_7_ (Table 1). On the dependence side, the majority of vitamin-dependent functions detected in the microbial community were ascribed to VB_1_, with 101,501 reads (0.78% of total reads) (Table 1), consistent with the importance of VB_1_-dependent enzymes in central metabolism (Table S1). VB_6_had the second highest number of vitamin-dependence transcripts (0.69% of all reads), followed by VB_7_ and VB_12_. Differences in the number of synthesis or dependence transcripts between the vitamins is likely to reflect variations in the complexity of their biosynthesis pathways, the number of reactions that require them, the times they can be reused, and the efficiency of their cellular salvage pathways.

**Table 1.**
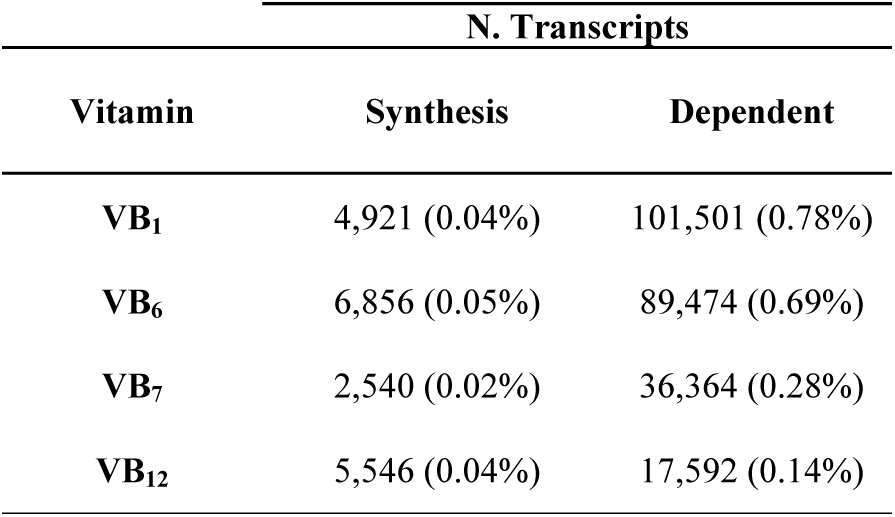
Total number of B-vitamin synthesis and dependence transcripts found in our study. Values in parentheses are the percentages of the B-vitamin reads in the dataset.

**Figure 2.**
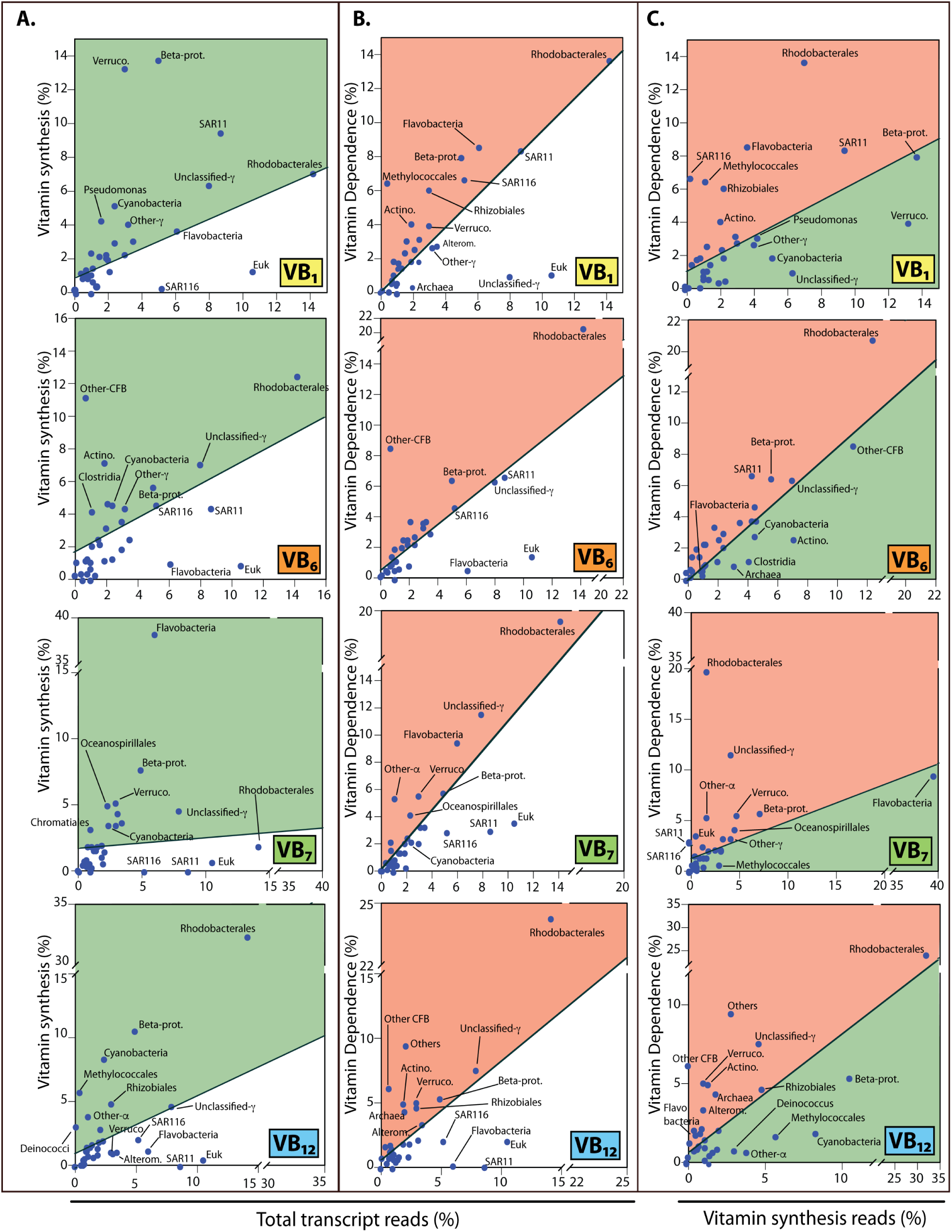
B-vitamin synthesis and dependence transcripts in different microbial taxa. A) Percentage of B-vitamin synthesis transcripts and B) percentage of vitamin dependent transcripts compared to the total transcript abundance for each taxonomic group. C) Percent-age of B-vitamin synthesis transcripts vs. percentage of B-vitamin dependence transcripts. Green areas show the microbial groups with a higher percentage of the community vitamin synthesis transcripts compared to total transcript abundance (A) or the percentage of their vitamin dependent gene transcripts (C). Red areas show the microbial groups with a higher percentage of vitamin-dependent transcripts as compared to total transcript abundance (B) or B-vitamin synthesis transcripts (C). It is assumed that the microbial groups on the regression line have a neutral impact in the community, as vitamin synthesis processes are relatively well balanced with their requirements. Regression lines are determined by outlier removal with IWLS (10^6^ maximum iterations, Table S4).

### Identification of B-vitamin synthesizers within the microbial community

We further analyzed the taxonomic distribution of vitamin synthesis and utilization transcripts among the members of the microbial community (Figure 2, Tables S3 and S2). Figure 2 shows the percent of community transcripts for vitamin synthesis or dependence that is attributed to a taxon. Some of the best-represented taxa dominated the expression of several vitamin synthesis pathways. Rhodobacterales, the most transcriptionally abundant bacterial group identified reached 7.5-10.5% of all transcripts during summer 08 and were also high but to a lesser extent in summer ‘09 (2-4%in this study, was potentially the major producer of VB^12,^ of all transcripts), while they were low in all other seasons (Hollibaugh et al., 2011; 2013). Other taxonomic groups such as Cyanobacteria and Verrucomicrobia were also present throughout the year although comprising less than 4% of the prokaryotic expression (Figure 1, Table S3).3 further shows the percentage of mRNA sequences that each phylogenetic group invested in B-vitamin synthesis and dependent processes. This analysis allows to qualitatively distinguish to what extent the different taxa rely on B-vitamin metabolism compared to the average community. Similarly to the total transcript analysis (Figure 2), Rhodobacterales invested a remarkably small percentage of their transcripts in VB_7_ synthesis, and to a lesser extent, in the production of VB_6_, and VB_1_. In contrast, the percentage of transcripts invested in VB_12_ synthesis and dependent processes by Rhodobacterales was close to the community average (Figure 3). This suggests that in order to maintain their high level of total expression in this microbial ecosystem (Figure 1), the Rhodobacterales may rely on other external vitamin sources (e.g. VB_7_ produced by other microbes).

**Figure 3.**
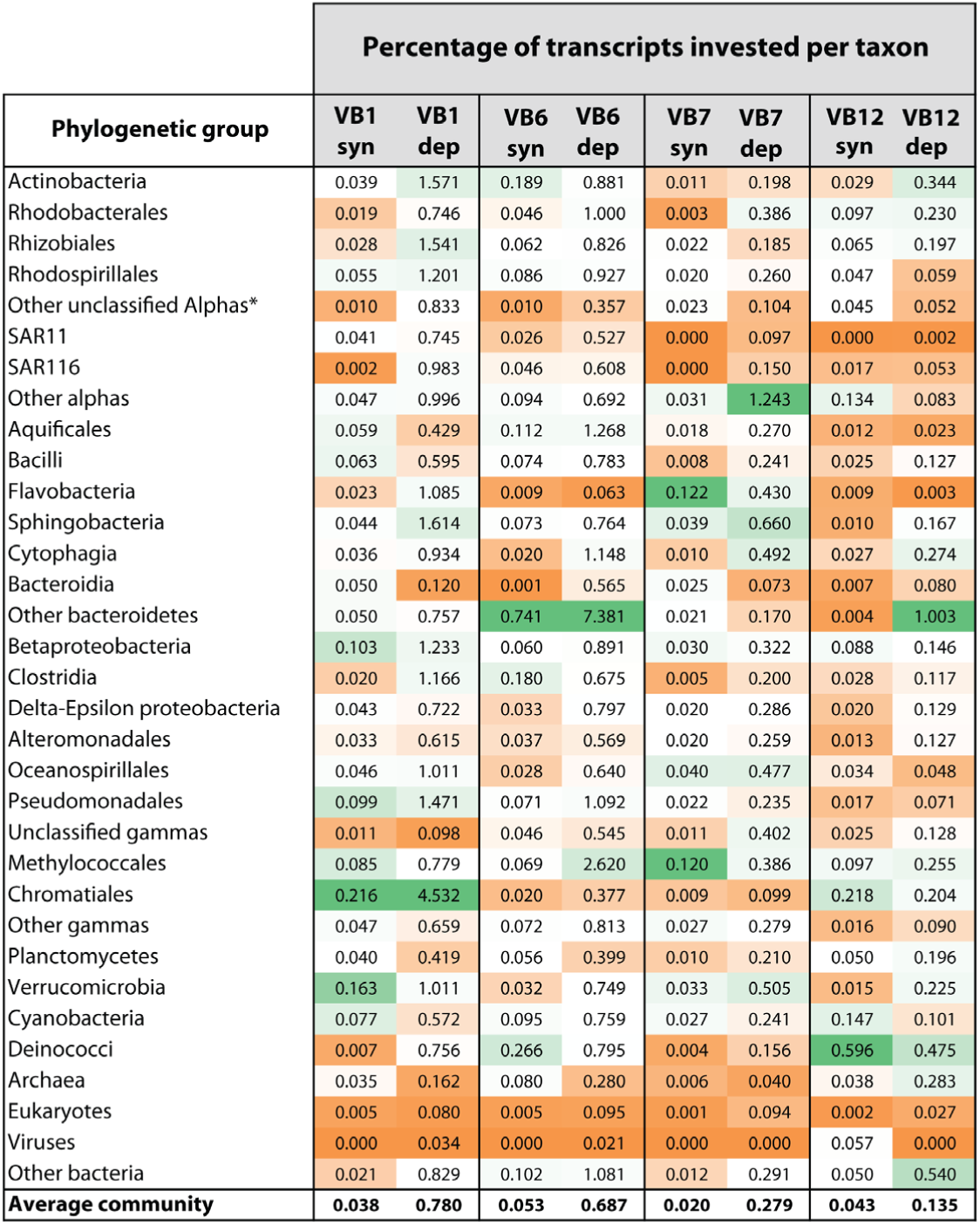
Percentage of transcripts invested in B-vitamin me-tabolism (B-vitamin synthesis or dependence) for each phylogenetic group. Heatmap colors designate the level of transcriptome investment compared to the average microbial community. White background represents the average percent investment, while green are above and orange are below average, respectively. *Un-classified alphaproteobacteria outside the SAR11 group.

Flavobacteria were also well represented and dominated the expression of VB_7_ synthesis genes, contributing as much as 64% of the total community synthesis transcripts for this vitamin (38% on average; Figure 2A, Table S3 and S4). Moreover, the VB_7_ requirements for this group were also higher than typical (Figure 2), suggesting a potentially high VB_7_ synthesis, usage and, thus, turnover for this group. It is currently known that different members of Flavobacteria, even close species of the same genus, have different vitamin-B_1_ synthesis capabilities (GÓmez-Consarnau et al., 2016). However, as a group compared to the community average, Flavobacteria expressed a higher percentage of VB_1_ dependence transcripts compared to synthesis (4% compared to 9% on average, respectively; Figure 2BC, Table S3 and S4). VB_12_ synthesis gene transcripts were low in Flavobacteria (1 % of the community). Furthermore, the expression of genes for VB_12_ dependent functions was almost absent (0.1% on average), suggesting a low requirement for this vitamin. This is consistent with the observation that none of the whole genome sequenced Flavobacteria available have a complete pathway for VB_12_ synthesis (Sañudo-Wilhelmy et al., 2014). Notably, the percentage of VB_7_ transcript synthesis was higher in this group than in any other taxon (0.12% of all Flavobacteria transcripts; Figure 3). The potentially significant VB_7_ contribution from Flavobacteria to the bacterial community and its VB_1_ dependence are also consistent with whole genome results showing that about 60% of the sequenced cultures of that group produce VB_7_ but only 25% can synthesize VB_1_ and none can make VB_12_ (Sañudo-Wilhelmy et al., 2014). However, our analysis showed that neither of the two Flavobacteria reference genomes that recruited the most transcripts in our study, Flavobacteria strains MS024-2A and MS024-3C, have the complete pathway to synthesize VB_7_ *de-novo* (Table S3). These organisms only partially expressed the VB_7_ synthesis pathway and would potentially require the combined enzymatic activity of other bacteria to be able to obtain VB_7_. Alternatively, it is possible that recruitment to Flavobacteria reference genomes split transcripts from a single population into different genome bins. Nevertheless, our data suggests that Flavobacteria, whether they have the complete gene set for VB_7_ or not, potentially occupy an important niche for its production in this community.

Members of the SAR11 and SAR116 clades were also well represented in our samples (9% and 5% on average; Figure 1, Table S3), and they are both auxotrophic for VB_1_ and VB_7_. However, *Candidatus* Puniceispirillum marinum IMCC1322, the SAR116 genome recruiting the greatest number of reads in our samples, can synthesize VB_12_, while members of the SAR11 clade neither synthesize nor have requirements for this vitamin (Giovannoni et al., 2005; Figure 2, Table S3). The percentage of community transcripts for both VB_12_ synthesis and dependence for SAR116 was the same (2 %), suggesting that their VB_12_ synthesis could meet their requirements (Table S3). SAR11 and SAR116 genomes have incomplete pathways for the synthesis of VB_1_ and can only produce one of the two moieties that form the thiamin molecule (4-methyl-5-(2-phosphoethyl)-thiazole, THZ) (Carini et al., 2014). SAR11 VB_1_ synthesis genes accounted for 10% of all the community transcripts while the VB_1_ dependent functions were about 8% (Figure 2). However, they would need to acquire the other VB_1_ moiety (4-amino-5-hydroxymethyl-2-methylpyrimidine, HMP) from the environment. In contrast to SAR11, SAR116 VB_1_ synthesis transcripts were nearly absent (0.3%) while their VB_1_-requiring transcripts made up to 7% of the community (VB_1_ dependence:synthesis residuals of 6%; Figure 2C, Table S3), suggesting a strong dependence on exogenous sources of VB_1_. In line with the total community transcriptomic data analysis (Figure 2), the percentage of SAR11 and SAR116 transcripts invested in B-vitamin metabolism was low, only being slightly above the average community value for VB_1_ synthesis in SAR11 and dependence in SAR116 (Figure 3).

Although the small eukaryotes (<3 µm) accounted for up to 38% of the total transcripts (Figure 1, Table S3), their B-vitamin synthesis genes always remained under 6% of the community B-vitamin synthesis transcripts (Figure 2A, Table S3). Furthermore, the eukaryotic B-vitamin dependent gene transcripts remained under 11%, suggesting that their B-vitamin requirements were also relatively low (Figure 2B, Table S3). This observation was consistent with the percent of eukaryotic transcripts invested in B-vitamin metabolism, which remained below the community average for all the vitamins (Figure 3). This was unexpected since eukaryotic phytoplankton have been considered major vitamin-dependent organisms of the microbial plankton (Croft et al., 2005; Tang et al., 2010). However, complex regulatory processes in response to iron limitation (Cohen et al., 2017) or riboswitch controls (McRose et al., 2014) may impact the expression of vitamin-related genes, as observed in eukaryotic alga. Another important consideration is that some eukaryotic phytoplankton could synthesize some vitamins using a non-canonical pathway. For instance, *Emiliania Huxleyi* can grow using either a complete VB_1_ molecule or only one of the VB_1_ –forming moieties (HMP) while lacking the genes that encode for the other moiety (HET-P; 4-methyl-5-b-hydroxyethylthiazole adenosine diphosphate), suggesting an alternative uncharacterized VB_1_ synthesis pathway (McRose et al., 2014). In contrast, picoeukaryotic algae (e.g. Micromonas and Ostreococcus) can utilize VB_1_ but not the individual moieties that form the vitamin (McRose et al., 2014, Paerl et al., 2015). Instead these picoeukaryotic phytoplankton are able to meet their VB_1_ requirements using a still chemically uncharacterized HET-P precursor together with HMP (Paerl et al., 2017). Finally, larger eukaryotic phytoplankton groups (>3 µm) were not included in this analysis and may have a more skewed ratio of synthesis to dependence transcripts.

In contrast to eukaryotic phytoplankton, betaproteobacterial B-vitamin synthesis genes were high for VB_1_ and VB_12_ (average: 14% and 10% respectively) and for VB_7_ (8%) (Figure 2; Table S3) and their vitamin dependence genes generally accounted for a similar percent of the community (close to the typical regression lines, Figure 2), suggesting that this bacterial group was very active and self-sufficient at producing and utilizing these B-vitamins. Consistent with this observation, the percentage of total betaproteobacterial transcripts invested in B-vitamin metabolism appeared to always be above the community average (Figure 3). However, there were significant differences in the number of VB_12_ synthesis and dependence transcripts within Betaproteobacteria (Table S2). While the abundant betaproteobacterial genomes dominated the VB_12_ requirement transcripts (e.g. Methylovorus glucosetrophus SIP3-4, beta proteobacterium KB13, or Methylophilales bacterium HTCC2181), the recruited reads for VB_12_ synthesis belonged to different Betaproteobacteria (e.g. Ralstonia solanacearum PSI07 and Chromobacterium violaceum ATCC 12472), which appeared to be much less abundant in this environment (Table S2).

Overall, our transcript data suggests that the vitamin synthesis and utilization potential is compartmentalized among the different members of the microbial community (Figures 1, 2, 3). Some microbial taxa representing a small percent of community transcripts appeared to be nonetheless important synthesizers of some B-vitamins (Figure 2A,Table S3), such as Verrucomicrobia with VB_1_ (<4% of total transcripts but 13% of VB_1_ synthesis transcripts; dependence:synthesis residual of -5%), Actinobacteria with VB_6_ (2% of total transcripts but 7% synthesis transcripts;dependence:synthesis residual of -4%), Cyanobacteria with VB_12_ (3% of total transcripts but 9% of synthesis transcripts; dependence:synthesis residual of -5%) and Methylococcales with VB_12_ (1% of total transcripts but 6% of synthesis transcripts; dependence:synthesis residual of -3%). Thus, less abundant members of the bacterial assemblage might also be important vitamin producers in the community, and therefore play larger ecological roles than their abundance would suggest. Our data suggest that this coastal marine microbial community is composed of organisms with distinct transcript contributions to vitamin synthesis and utilization. These expression patterns are certainly complex and may be the result of a combination of diverse regulatory processes that are specific to some taxonomical groups (e.g. eukaryotic phytoplankton; McRose et al., 2014) and are difficult to discern at the community level. Future controlled laboratory studies using specific members of the microbial community will be needed to remove any constraints in our field results.

### Seasonal and circadian regulation of B-vitamins gene expression and their potential impact on water-column dissolved vitamin concentrations

We hypothesized that the ambient dissolved vitamin concentrations measured in the water column should reflect, to some extent, a balance between the synthesis and utilization by the microbial community. During our study, the concentrations of dissolved VB_7_ and VB_12_ were the lowest of all, and below our detection limits. These low levels of VB_7_ and VB_12_ might suggest a very tight synchronization between synthesis and utilization that may not allow any build-up of these two vitamins in this coastal environment. This hypothetical tight synthesis-uptake coupling, however, was not observed at the level of gene expression, as the dependence:synthesis transcripts for VB_12_ and VB_7_ varied over time with no clear temporal patterns (Figure 4). In this study we only quantified the cyanocobalamin form of VB_12_ and we cannot rule out the presence of the other chemical forms of this vitamin (e.g., pseudo-methyl-, adenosyl-and hydroxo-cobalamin; Suarez-Suarez et al., 2011; Helliwell et al., 2016; Heal et al., 2017; Suffridge et al., 2017). It is unclear how the availability of the different chemical forms of VB_12_ relate to gene expression and future studies will need to address that issue

**Figure 4.**
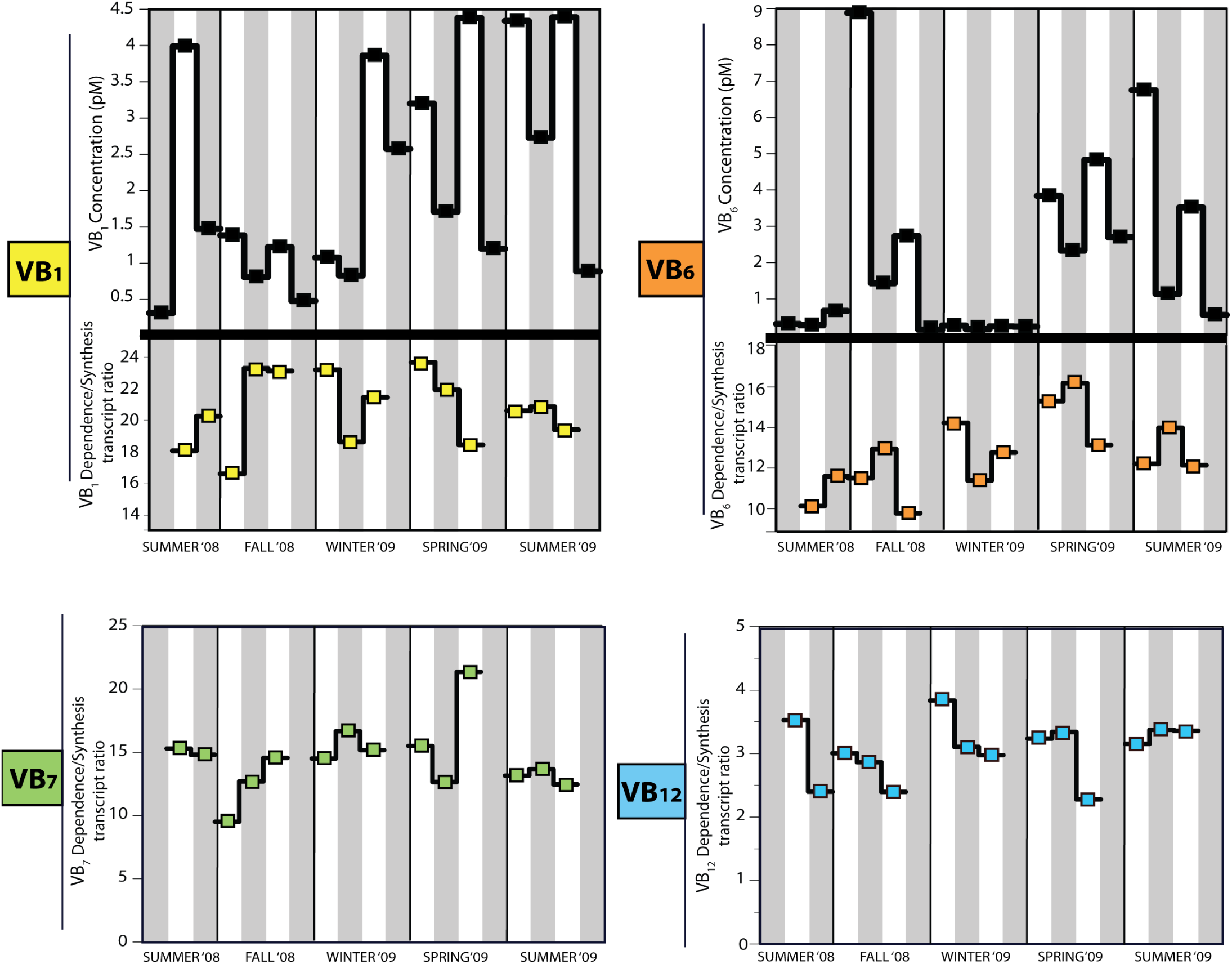
Environmental concentrations of dissolved VB_1_ and VB_6_ compared to the vitamin dependence:synthesis transcript ratios. Concentrations of dissolved VB_7_ and VB_12_ were below our detection limit and only the vitamin dependence:synthesis transcript ratios are shown. Grey and white background denotes night and day samples, respectively. Specific concentrations and times of sampling are shown in Table S6.

Dissolved VB_1_ ranged from 0.1 to 5 pM and followed diel oscillations, with consistently higher levels during the day compared to night within each season (Figure 4). However, as for VB_7_ and VB_12_, the VB_1_ dependence:synthesis transcripts did not show any clear diel or seasonal dynamics (Figure 4). We attributed the higher dissolved VB_1_ concentration during only meet their biological demands with the uptake of VB_1_ precursors instead of the complete VB_1_ molecule (Carini et al., 2014). Even though we did not quantify the different moieties of this vitamin, knowing the standing stock of VB_1_ the day to bacterioplankton VB_1_ production or excretion, as most of the synthesis transcripts belonged to heterotrophic bacteria rather than to small phytoplankton. (Figure 2A,Table S3). Nonetheless, we cannot rule out the impact of the large phytoplankton (>3µm) on vitamin levels as they were not included in our samples. One explanation for the overall higher VB_1_ concentrations in the day samples could be its function as photo-protectant to oxidative stress during light exposure, as shown for Escherichia coli and Arabidopsis thaliana (Jung and Kim, 2003; Tunc-Ozdemir et al., 2009). Notably, Gifford et al. (2014) showed that the activity of Rhodobacterales (inferred by the expression of ribosomal proteins) in the same metatranscriptomes was also influenced by diel cycles. These similar patterns could indicate a relationship between the activity of Rhodobacterales and the observed higher concentrations of dissolved VB_1_ found in the day samples, as most Rhodobacterales are auxotrophic for this vitamin or at least for one of its moieties (Table 2). However, the matter of VB_1_ availability is further is relevant, as some of its decomposition products could be used for cellular growth by diverse phytoplankton groups (Gutowska et al., 2017). Future studies will need to address the effect of VB_1_ moieties and pathway intermediates in natural communities as recently established in culture growth studies (Gutowska et al., 2017; Paerl et al., 2017).

**Table 2.**
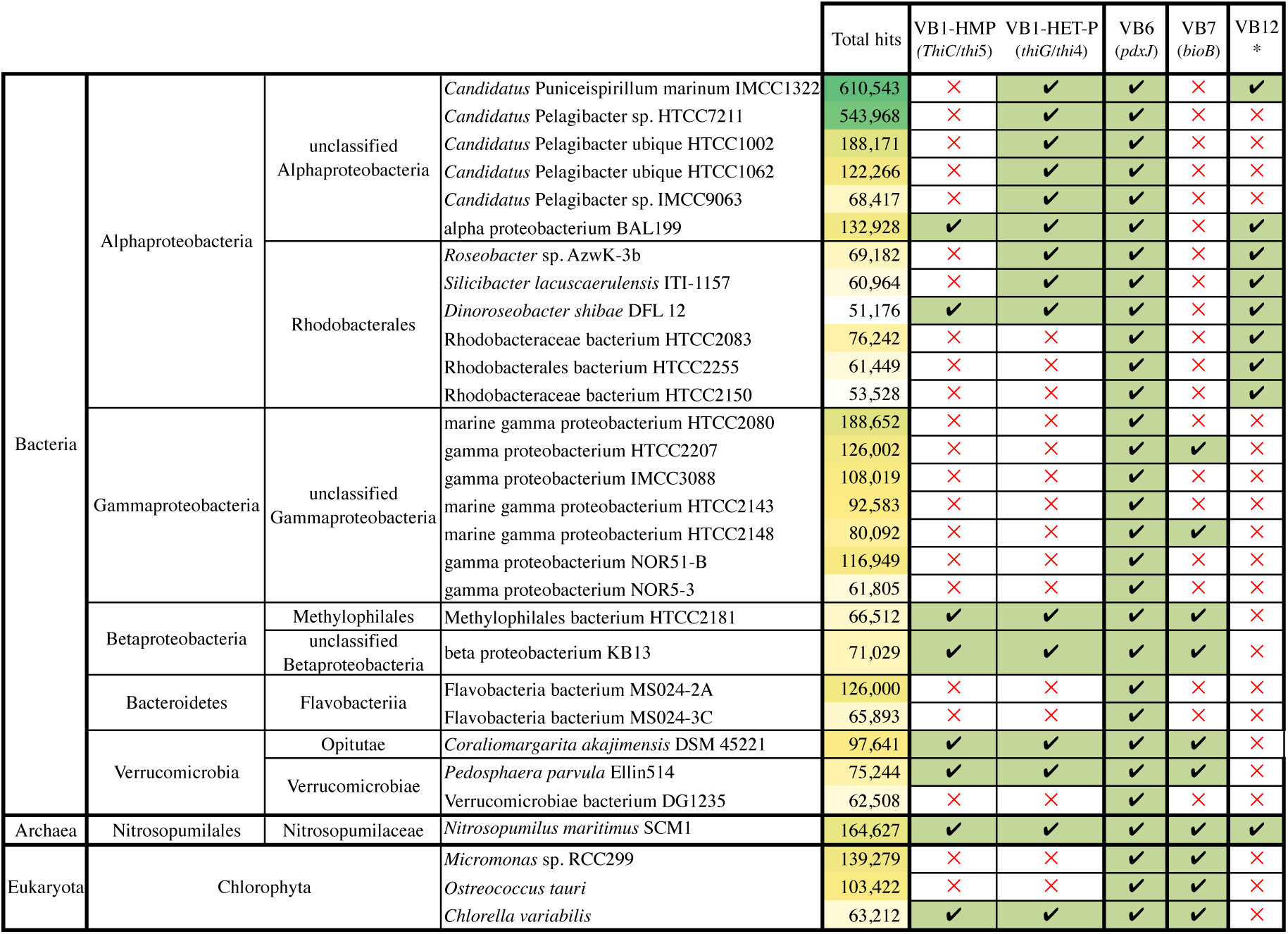
Presence of B-vitamin synthesis genes and pathways for the 30 most abundant genomes (according to total transcripts) found in the Sapelo island metatranscriptome. Vitamin-B_1_ synthesis was determined using the thiC-thi5 and thiG-thi4 genes for the production of the 4-amino-5-hydroxymethylpyrimidine (HMP) and 4-methyl-(β-hydroxyethyl) thiazole (THZ) moieties, respectively; Vitamin-B_6_ synthesis was determined by the presence of the pdxJ gene. Vitamin-B_7_ synthesis was determined using the bioB biotin synthase gene. To account for possible misannotations of single genes on the vitamin-B_12_ synthesis pathway, it was considered pres-ent when a genome contained more than 75% of the VB_12_ *de-novo* synthesis clusters of orthologous groups (COGs) retrieved from the Integrated Microbial Genomes database (http://img.jgi.doe.gov) (COG0007, COG0310, COG0368, COG1010, COG1270, COG1429, COG1492, COG1797, COG1903, COG2073, COG2082, COG2087, COG2099, COG2109, COG2241, COG2242,and COG2243), as previously reported by Sañudo-Wilhelmy et al. (2014).

The levels of dissolved VB_6_ ranged from 0.1 to 8.9 pM and were also higher in the day samples, except for winter ‘09 when all VB_6_ concentrations were below our detection limit (<0.1 pM; Figure 4). Similar to VB_1_, the higher VB_6_ concentrations during the day could be explained by its function as antioxidant during light exposure (Bilski et al., 2000; Mooney et al., 2009). In fact, the antioxidant properties of VB_6_ against oxidative stress even exceed those of vitamins C and E (Ehrenshaft et al., 1999). Furthermore, this coenzyme is particularly important in amino acid metabolism (e.g. amino acid synthesis and transaminations; Hayashi, 1995), which may be more relevant during the night (Cuhel et al., 1984; Poretsky et al., 2009; Gifford et complicated by the fact that several microbial groups can al., 2014). Yet, we did not observe diel oscillations in VB_6_ dependence:synthesis transcript ratios.. This is in contrast to a metatranscriptomic study in the South Pacific, in which the expression of VB_6_ synthesis genes was higher during the night compared to the day (Poretsky et al., 2009). Additional studies analyzing intracellular B-vitamin concentrations will be necessary to identify their circadian fluctuations in relation to the internal cellular processes over time.

The observed higher dissolved B-vitamin concentrations during sunlight exposure could have important consequences for the functioning of the marine microbial community. An obvious one is that vitamin auxotrophs that take advantage of this source would end up following the same circadian rhythm as the vitamin producers. Ottesen et al. (2014) reported a tight diel synchronization between the metatranscriptomes of the cyanobacteria Prochlorococcus and heterotrophic bacterioplankton in situ, suggesting an interaction (direct or indirect) between these organisms (Ottesen et al., 2014; Armbrust, 2014). In our study, dissolved VB_1_ concentrations were positively correlated to Chl-a throughout the yearlong dataset (Figure 5). Although a direct cause-effect relationship between the two parameters has not been demonstrated, this vitamin is involved in carbon fixation during the Calvin cycle (Sañudo-Wilhelmy et al., 2014; Monteverde et al., 2017). Because the VB_1_ dependent transcripts of eukaryotic picophytoplankton were low, the correlation between dissolved VB_1_ and total Chl-a may involve the larger phytoplankton community not sampled by our size-fractionation scheme (0.2 > 3 µm). Further studies will be needed to evaluate the cause of the observed VB_1_/Chl-*a* relationship and confirm the identity of the key organisms responsible for this interdependence. In contrast to the strong correlation between VB_1_ and Chl-a,we did not find any relationship between the concentrations of dissolved VB_6_ and phytoplankton biomass (Figure 5).

**Figure 5.**
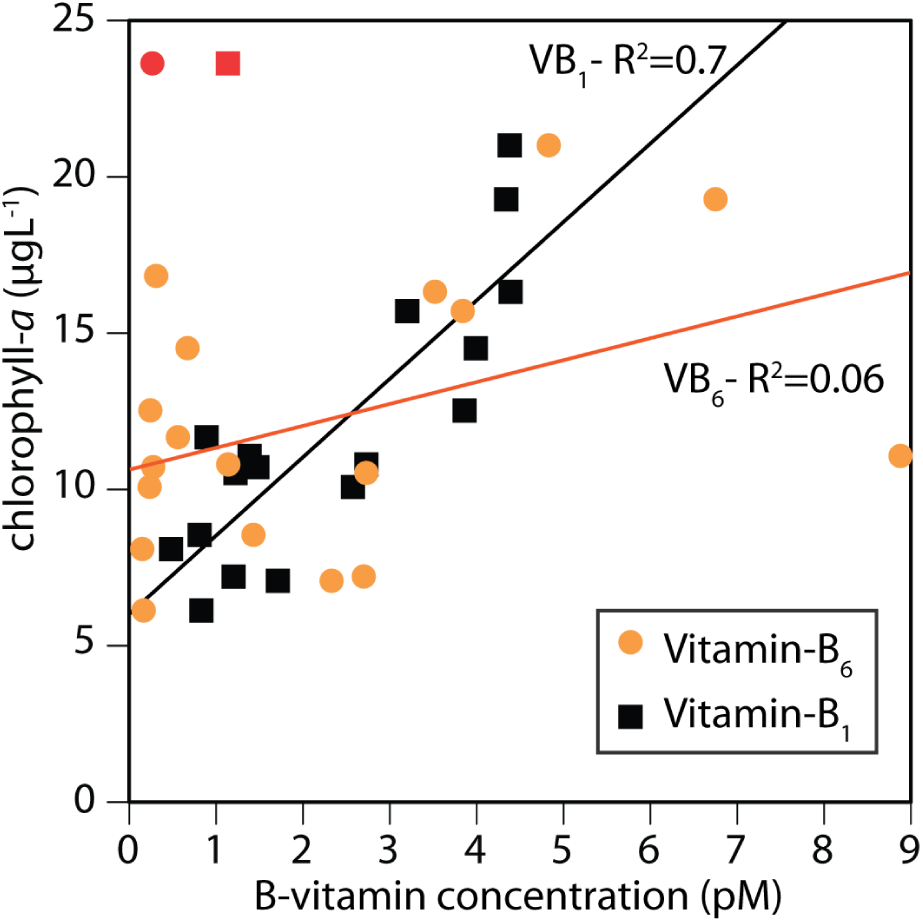
Correlation between total chlorophyll-a and dissolved VB_1_ and VB_6_ concentrations. The red data points are considered outliers and were excluded from the statistical analysis; those correspond to the sample collected in winter ‘09.

Although the auxotrophy for VB_1_, VB_7_ and VB_12_ has been recognized in marine microbes for decades (Provasoli, 1963; Provasoli and Carlucci, 1974; Croft et al., 2005; Sañudo-Wilhelmy et al., 2014), VB_6_ dependence had not yet been considered. Indeed, VB_6_ auxotrophy is likely to be minor as the VB_6_ synthesis pathway is widespread in free-living microorganisms and plants and more than a hundred enzymatic reactions are catalyzed by this coenzyme (Hayashi, 1995). Nonetheless, our results suggest that some members of the microbial community express more VB_6_ synthesis transcripts than others in relation to their total transcript contribution to the community and their vitamin requirement transcripts (Figure 2). Moreover, if the excess of VB_6_ is released to the environment on daily basis, it is possible that at least some taxa have adapted to take advantage of this source to reduce their vitamin synthesis requirements. Finally, because the inferred vitamin metabolism in this coastal community involves bacterial groups that are cosmopolitan, our findings could potentially be extrapolated to other marine environments.

## Experimental procedures

### Sample collection

Samples for metatranscriptomics were collected quarterly (2008: August 6–7, November 5–7; 2009: February 15–17, May 13–15, August 12–14) at Marsh Landing, Sapelo Island, Georgia, U.S.A. (31°25’4.08 N, 81°17’43.26 W) as part of the Sapelo Island Microbial Observatory program (http://simco.uga.edu). Sampling occurred at four consecutive high tides resulting in two consecutive pairs of day-night samples per season. Additional samples were collected for specific environmental measurements including chlorophyll-a and dissolved B-vitamins. RNA from the water samples was extracted as previously described (Poretsky et al., 2009; Gifford et al., 2011; 2013; 2014). Briefly, 6-8 L were directly filtered for 11-14 min from a depth of 1 m through a 3-µm pore-size prefilter to exclude larger microbial fractions (Capsule Pleated Versapor Membrane; Pall Life Sciences, Ann Arbor, MI, USA) and 0.22-µm pore-size filter (Supor polyethersulfone; Pall Life Sciences). The 0.22-µm filter was immediately flash frozen in liquid nitrogen until RNA processing and sequencing.

### RNA processing and sequencing

Samples were processed as earlier described (Gifford et al., 2011; 2013; 2014). Total RNA was extracted using a RNAEasy kit (Qiagen) and DNA was removed using a Turbo DNA-free kit (Applied Biosystems, Austin, TX, USA). mRNA was enriched via enzymatic rRNA reduction using Epicentre’s mRNAOnly isolation kit (Madison,WI, USA) and subsequently with MICROBExpress and MICROBEnrich rRNA (Applied Biosystems) subtractive kits. Enriched mRNA samples were linearly amplified using the MessageAmp II-Bacteria kit (Applied Biosystems) and double stranded cDNA synthesized with Promega’s Universal RioboClone cDNA synthesis system and random primers. cDNA synthesis reactions were cleaned up with a QIAquick PCR purification kit (Qiagen). Samples were sheared and size-selected to ∼300bp and sequenced with an Illumina GAIIX to obtain 150×150 bp paired end reads.

### Bioinformatics pipeline

Reads with an average quality score ≤ 20 and a length <100 bp were removed and overlapping paired end reads from each fragment were assembled using SHERA (Rodrigue et al., 2010) with a score >0.5 (Gifford et al., 2013; 2014). Sequences were combined from all samples, subsampled to 25,000 reads, and searched against the SILVA small and large subunit databases using BLASTn (Altschul et al., 1998) with bit score ≥ 50 to create a more compact database for further rRNA gene filtering. The total reads from all metatranscriptomes were then filtered using BLASTn against the compact database. Putative non-rRNA sequences were then searched against the National Center for Biotechnology Information’s (NCBI; http://www.ncbi.nlm.nih.gov) RefSeq protein database (version 43, September 2010) using BLASTx with a bit score cutoff ≥ 50 to identify protein encoding sequences. Functional annotation and taxonomy were assigned based on the top scoring hit to RefSeq. RefSeq counts for each sample were randomly subsampled (van Rossum, 2010; McKinney, 2010) to the total number of non-rRNA reads of the smallest library (1,332,199 reads) to account for variation in sample sequencing effort.

### Annotation of vitamin related transcripts

Vitamin related transcripts for VB_1_ (thiamin monophosphate; TMP or thiamin pyrophosphate; TPP), VB_6_ (pyridoxal 5’-phosphate), VB_7_ and VB_12_ (methylcobalamin, adenosylcobalamin, hydroxocobalamin) were identified by text-based query for either the EC number, gene or enzyme name in the retrieved RefSeq functional annotations. The specific vitamin synthesis genes and vitamin dependent enzymes are listed in Table S1 and all vitamin-related hits are included in supplementary Table S2. For each of the vitamins (VB_1_, VB_6_, VB_7_, VB_12_), vitamin dependent enzymes were identified using the ExPASy database (http://www.expasy.org) and are defined as the metabolic reactions that require a specific B-vitamin as co-enzyme. Vitamin synthesis functional annotations are those enzymes that belong to the synthesis pathway of a specific B-vitamin, and were determined using the Kyoto Encyclopedia of Genes and Genomes (KEGG; http://www.genome.jp/kegg/)(Table S2). For VB_7_ synthesis pathway, the genes fabI, fabB, fabG and fabZ were not included in our analysis, as they are also part of the fatty acid biosynthesis pathways and are present in most microorganisms including vitamin-B_7_ auxotrophs. Therefore includingthose geneswouldproducefalsepositives. Sequences were also classified by sequence similarity to a list of vitamin synthesis/dependence genes in order to verify that RefSeq description annotation precisely represents counts of vitamin synthesis and dependence. Sequences for synthesis and dependence of each vitamin were retrieved from the RefSeq protein database. All sequences were searched against separate BLAST databases for synthesis and dependence of each vitamin (e. g. VB_6_ dependence and VB_6_ synthesis). Only matches with a bit score ≥ 40 and that do not have a higher scoring non-vitamin RefSeq match were retained. This strategy produced slightly underestimated counts of vitamin synthesis/dependence classifications (86 ∓ 4.2%; Table S5), but did not affect the overall pattern of vitamin usage.

### Identification of taxa involved in B-vitamin synthesis and utilization

Linear modeling robust to outliers was employed to identify taxa that disproportionately contributed to vitamin synthesis or dependence gene expression above the typical expression for the whole community. Iterated re-weighted least squares (IWLS) linear regression was fit with 1 x106 maximum iterations for outlier removal between synthesis:total expression, dependence:total expression, and dependence:synthesis using the “MASS” R-package (Venables and Ripley, 2013). Ratios near the linear regression line (low residuals) are defined as typical for the microbial community. A positive residual suggests a ratio greater than the typical ratio, whereas a negative residual is less than the typical ratio. For example, the negative Cyanobacteria dependence:synthesis residual for VB_1_ suggests that Cyanobacteria are important synthesizers of VB_1_. Underlying this analytical approach is the assumption that vitamin requirements scale similarly with vitamin-dependent transcripts for all taxonomic groups (i.e., a vitamin dependent transcript represents an equivalent vitamin requirement across all taxa). The synthesis:dependence regression lines of each vitamin could be influenced by the fact that particular taxa are auxotrophs for a particular vitamin (e.g. eukaryotes and VB_12_), and by different taxa having varied numbers of enzymes that require the vitamin. For instance, there are more than 20 enzymes that require VB_12_ in prokaryotes (Marsh 1999), and only three VB_12_ dependent enzymes in eukaryotes (Helliwell et al., 2011). Another example is VB_1_ which, among other functions, is used by enzymes in the Calvin cycle of photoautotrophs and this cycle will be absent in heterotrophic microorganisms. Finally, as in any other metatranscriptomic analysis, our results could not account for any post-transcriptional regulation.

### Chlorophyll-*a* quantification

Chlorophyll a analysis was done as described in Parsons et al.(1984). Seawater samples of 250 ml were filtered onto GFF membranes (Whatman plc, Maidstone, United Kingdom), placed in conical tubes filled with 50 ml of 90% acetone, and stored at -20°C until analysis. Sample fluorescence was measured by sonicating the samples for 30 seconds, then adding 3 ml to a 5 cm quartz cuvette, and measuring on a Turner fluorometer. Estimates of phaoe-pigments were obtained by acidifying the same sample with 10% HCl and rerunning on the fluorometer. Sample chlorophyll-a concentrations were then calculated based on comparison to a standard curve of fluorescence from chlorophyll-a standards.

### Quantification of dissolved B-vitamins in situ

The concentration of dissolved B-vitamins was determined using HPLC-MS after pre-concentration of 500 mL of 0.2 µm-filtered seawater. Dissolved samples were preconcentrated by passing the sample over a C18 resin at two pHs (6.5 and 2.0), followed by elution with 12 ml of methanol. A nitrogen (N_2_) dryer was used to evaporate the samples to about 250 µl. Quantification of dissolved B-vitamins, VB_1_ as thiamin hydrochloride (C_12_H_17_ClN_4_OS*HCl), VB_6_ as pyridoxine hydrochloride (C_8_H_11_NO_3_*HCl), VB_7_ asbiotin(C_10_H_16_N_2_O_3_S) and VB_12_ as cyanocobalamin (C_63_H_88_CoN_14_O_14_P), was conducted using a Thermo Scientific Quantum Access electrospray ionization triple quadrupole mass spectrometer, coupled to a Thermo Scientific Accela High Speed Liquid Chromatography (LC/MS) system. The LC system used a stable-bond C18 reversed-phase column (Discovery HS C18 10 cm×2.1mm, 5µm column, Supelco Analytical), with a methanol:water gradient program. A full description of the analytical protocol for the vitamins quantification including all of the MS operating conditions have been reported elsewhere (Sañudo-Wilhelmy et al., 2012). Although our analytical protocol is able to measure a suite of B-vitamins, the small volume of the water samples did only allow the determination of vitamins found at relatively high concentrations such as thiamin and pyridoxine (both measured against standards prepared in their hydrochloride forms). The concentrations of dissolved VB_7_ and VB_12_ were below our detection limit in all samples (<0.1 pM).

## Acknowledgments

This project was founded by the Marie Curie Actions–International Outgoing Fellowships (project 253970) and the US National Science Foundation grant OCE-1435666.

## Author contributions

LG-C and MM designed the study. SG collected and processed field samples. LC and SS-W analyzed B-vitamin concentrations. LG-C, RS, SG, LC, JF,SS-W, MM analyzed data and contributed to the writing of the paper.

## Author information

The authors declare no conflict of interest. Correspondence and request for materials should be addressed to LG-C.

## Supporting Information

Additional Supporting Information may be found in the online version of this article:

**Table S1.** List of B-vitamin synthesis and B-vitamin dependent functions used in the metatranscriptome analysis. Vitamin dependent functions were identified using the ExPASy database (http://www.expasy.org) and vitamin synthesis genes were identified using the Kyoto Encyclopedia of Genes and Genomes (KEGG; http://www.genome.jp/kegg/). In addition to the listed enzymes, the reads annotated as “vitamin synthesis” or “vitamin dependent/requiring” included in Table S2 and Table S3 were also included.

**Table S1.** List of all B-vitamin synthesis and B-vitamin dependentfunctionreadsidentifiedinthemetatranscriptomics dataset. In addition to the enzymes listed in table S1, reads annotated as “vitamin synthesis” or “vitamin dependent/ requiring” were also included.

**Table S1.** Summary of the B-vitamin synthesis and dependent gene transcript abundance for the best-represented taxonomic groups in each individual sample.

**Table S1.** Percentages of B-vitamin synthesis and dependence transcripts in different microbial taxa and regression lines shown in Figure 2.

**Table S1.** Comparison of annotation methods for vitamin synthesis and dependence transcripts. To verify that RefSeq description annotation accurately represents counts of vitamin synthesis and dependence, reads were also classified by sequence similarity to a list of vitamin synthesis and dependence genes. Reference sequences for synthesis and dependence genes of each vitamin were retrieved from the RefSeq protein database. Metagenomic reads were searched against separate BLAST databases for the synthesis and dependence genes associated with each vitamin. Only matches with a bit score >= 40 and that did not have a higher scoring non-vitamin RefSeq match were retained.

**Table S1.** Environmental concentrations of Chlorophyll-a, vitamin B_1_ and B_6_ and specific sampling times.

